# MAT-classifier: A memory-efficient pipeline for accurate genus-level profiling from ancient metagenomic data

**DOI:** 10.64898/2026.01.28.702372

**Authors:** Aranya Dhibar, Mikhail V Matz

**Affiliations:** Department of Integrative Biology, University of Texas at Austin, Austin, Texas, USA

**Keywords:** ancient DNA, ancient metagenomics, ancient microbiome, taxonomic classification, memory-efficient pipeline, accurate profiling

## Abstract

**Background:** Advances in sequencing technology have expanded opportunities to recover microbial DNA from ancient samples and reconstruct past environments and host–microbe interactions. However, the field remains constrained by computational challenges and accuracy problems, as rare ancient microbial DNA must be distinguished from abundant modern contaminants. Moreover, existing pipelines demand substantial computational resources, particularly memory, limiting their accessibility.

**Results:** Here, we present MAT-classifier, a genus-level profiling workflow for detecting ancient microbial taxa from metagenomic projects, designed to reduce computational requirements while increasing accuracy. Unlike its counterparts, MAT-classifier first consolidates candidate references at the genus level and then performs independent alignments using conventional short-read aligners instead of metagenomic aligners. Using simulated datasets, we showed that this approach achieves more accurate classification of ancient taxa while requiring substantially less memory and shorter runtime than a modern counterpart, the aMeta pipeline. Benchmarking on multiple empirical ancient datasets further confirmed its low memory footprint and practical utility.

**Conclusions:** MAT-classifier provides a reliable, computationally efficient, and accessible framework for ancient microbiome profiling. It lowers computational barriers while maintaining robust classification performance, facilitating broader application of ancient microbial DNA analysis.

## Background

The field of ancient DNA (aDNA) research has traditionally centered on evolutionary questions involving humans and other fauna, restricting the focus to eukaryotic samples. Advances in next-generation sequencing have expanded the scope by enabling extensive sequencing of ancient samples at an increasingly lower cost. High-depth metagenomic sequencing has opened the avenue to study the host-associated microbial aDNA preserved within eukaryotic remains, as they often constitute an abundant non-host component alongside modern contaminants. Ancient microbial DNA can provide valuable information about the past, for example, ancient pandemics, lifestyle, and population migrations (Mühlemann et al., 2018; Rascovan et al., 2019; Rasmussen et al., 2015; Spyrou et al., 2019). In parallel, the emergence of sedimentary ancient DNA (sedaDNA) as an independent branch of palaeogenetics has further highlighted the importance of ancient microbes alongside other organisms (Gelabert et al., 2021; Pedersen et al., 2021; Slon et al., 2017; Vernot et al., 2021; Wang et al., 2021; Zavala et al., 2021).

Despite its promise, even with extensive shotgun sequencing (Orlando et al., 2021) ancient microbiome research often suffers from challenges in taxonomic classification and authentication. Ancient microbial sequences are often short, damaged, and rare within a complex background of modern environmental contaminant DNA, resulting in detection errors and authentication errors (Campana et al., 2014; Der Sarkissian et al., 2021). Detection errors occur when true taxa are missed, or reads are incorrectly classified, whereas authentication errors, an extra requirement for ancient DNA classifiers, lead to modern contaminants being mislabeled as ancient taxa. To decrease these errors, existing ancient taxonomic classification workflows rely on alignment-based methods, like the metagenomic aligner MALT (Herbig et al., 2016), as these alignments can be evaluated for post-mortem DNA damage patterns (5’ C>T and 3’ G>A substitutions - a signature of ancient DNA) using tools such as PMDtools (Skoglund et al., 2014) or mapdamage (Jónsson et al., 2013). However, the computational demand of aligning reads against large metagenomic reference databases presents a new challenge.

Several existing workflows combine metagenomic classification with downstream authentication for ancient DNA, including nf-core/mag (Krakau et al., 2022), HOPS (Hübler et al., 2019), and aMeta (Pochon et al., 2023), each with its own strengths and weaknesses. The nf-core/mag pipeline assembles metagenome-assembled genomes (MAGs) and authenticates ancient MAGs after de-novo assembly. However, ancient DNA studies rarely recover enough ancient reads to assemble MAGs, limiting applicability.

Assembly-free approaches such as HOPS use MALT to align reads against reference databases and then perform multiple downstream authenticity checks. More recent workflows, including aMeta, use a hybrid strategy in which ultra-fast k-mer-based classifiers first identify candidate taxa and reduce the reference database size before MALT alignment (Pochon et al., 2023; Ravishankar et al., 2025). Nevertheless, the competitive alignment performed by MALT remains computationally demanding because the whole reference database must be loaded into memory. This requirement increases memory usage and limits the analysis of large ancient metagenomes on modest computational systems. For instance, even after reducing the MALT database, aMeta recommends 256 GB of RAM for small projects, with requirements increasing for deeper sequencing projects (Eisenhofer & Weyrich, 2019; Pochon et al., 2023).

To address this limitation, we developed the **Metagenomic Ancient Taxa Classifier (MAT-classifier)**, a computationally efficient workflow that classifies ancient taxa with a non-competitive alignment approach along with genus-level reference binning and multi-metric authentication. MAT-classifier retains the initial taxonomic screening strategy used by previous hybrid workflows to identify candidate reference taxa using Kraken2 (Wood et al., 2019). But its principal methodological difference is that it does not perform the large-scale competitive alignment against all candidate genomes. Instead, MAT-classifier groups reference genomes by genus and aligns reads independently against each genus-level reference set using traditional aligners like Bowtie2 (Langmead & Salzberg, 2012) or BWA-aln (Li & Durbin, 2009). This design reduces memory requirements while parallel execution across genera improves computational efficiency.

Genus-level binning also addresses an important limitation of the independent mapping approach. Closely related species can produce ambiguous alignments, making species-level assignments particularly unreliable with short ancient reads. By grouping related reference genomes within genera, MAT-classifier avoids overinterpreting species-level differences, reduces redundant mapping among closely related species, and pools evidence across species. The resulting alignments are then evaluated using multiple authentication checks, including post-mortem damage estimated with PMDtools (Skoglund et al., 2014) and pyDamage (Borry et al., 2021), short fragment length, edit distance, and coverage evenness. These metrics are integrated into a scoring framework that enables easier identification and interpretation of candidate ancient taxa. Thus, MAT-classifier is designed to be a practical alternative to memory-intensive pipelines while preserving alignment-based evidence required for robust ancient microbial DNA authentication.

## Methods

### Kraken-based filtering and reference database construction

To keep the database size manageable (and reduce the number of alignment jobs), we implemented a Kraken2-based identification and filtering step to select candidate taxa. For this step, MAT-classifier requires Kraken2 reports from which it can dynamically construct a reference database. Kraken2 classification was kept outside the automated workflow because it is the most memory-intensive step (Supplementary Figure 1), allowing users to optimize Kraken2 execution for their available resources. We chose Kraken2 as it requires much less memory compared to KrakenUniq for the same database type (Langmead, 2025), but users can use KrakenUniq reports as well.

Kraken2 must be executed with the *--report-minimizer-data* option, which reports the number of distinct minimizers associated with each taxon. Minimizers are subsequences derived from k-mers; more minimizers indicate that classifications are supported across more independent regions of the reference sequence rather than concentrating in the same region. Read counts and distinct minimizers were used as proxies for depth and breadth of taxonomic support, respectively. Candidate species were selected if they have ≥100 reads and ≥500 distinct minimizers to support; genus-level classifications with >200 reads were also retained. These thresholds were found to be the best from simulations to balance false-positive and false-negative classifications (Supplementary Table 1).

For retained taxa, MAT-classifier retrieves genome assemblies from NCBI using the Datasets command-line interface, prioritizing the latest reference genomes and otherwise selecting an available assembly of at least contig-level quality. Genomes belonging to the same genus are concatenated into a single FASTA reference, producing genus-level reference sets for downstream alignment.

### Bowtie2 and BWA-aln alignment

Reads are independently aligned against each genus-level reference using either Bowtie2 (Langmead & Salzberg, 2012) or BWA-aln (Li & Durbin, 2009). Bowtie2 was run in “end-to-end” mode using the “--very-sensitive” preset, whereas BWA-aln was run using parameters commonly applied to short ancient DNA (-l 16500 -n 0.01 -o 2; (Dolenz et al., 2024; Green et al., 2010)). We test both these aligners with our pipeline. Bowtie2 performs better for USER-treated ancient DNA (Poullet & Orlando, 2020) but may exhibit higher reference bias (Dolenz et al., 2024; Oliva et al., 2021). On the other hand, BWA-aln may show lower levels of reference bias for shorter reads (Dolenz et al., 2024). Users can select either aligner.

Independent genus-level alignments are executed in parallel, with four threads assigned to each alignment job and the number of parallel jobs determined by user-specified CPU resources. Resulting alignments are deduplicated and sorted before downstream analysis.

### Validation, Authentication, and Scoring

Candidate genera are evaluated using five scoring metrics spanning four types of evidence: post-mortem damage, fragment length, taxonomic support, and edit-distance profiles.

Post-mortem damage is assessed using pyDamage (Borry et al., 2021) and PMDtools (Skoglund et al., 2014). pyDamage evaluates whether terminal damage significantly differs from its null model, with *q-value* < 0.05 considered evidence of ancient DNA. PMDtools evaluates damage at the individual-read level and assigns likelihood scores. A genus is supported if it has more than 5% of reads passing the PMD score threshold of 3. Users can change the percentage of reads passing the threshold.

Fragment length provides an additional authentication metric because degraded ancient DNA is generally shorter than modern DNA. MAT-classifier calculates the mean length of aligned reads per genus and evaluates the variation in mean read length across all genera within a sample. When read-length distribution has an interquartile range greater than 3 base pairs, k-means clustering (*k* = 2) is applied to separate read lengths into shorter and longer length clusters. If mean read lengths of genera fall within the shorter length cluster, they receive support for ancient origin.

Taxonomic presence is further validated using the Kraken2 report. Genera are supported and scored when they have >200 reads and >1,000 distinct minimizers, representing sufficient depth and breadth of classification. Alignment quality is independently evaluated from the edit-distance distribution. Support is assigned when most of the reads have zero edit distances, i.e., no mismatches, and the number decreases with increasing numbers of edit distance, i.e., mismatches.

MAT-classifier integrates these supports into a five-point scoring system, with one point assigned for each of the following: (i) significant pyDamage support (*q* < 0.05 and predicted accuracy > 40%), (ii) more than 5% reads have PMD >3, (iii) membership in the shorter read-length cluster, (iv) sufficient Kraken2 read and distinct-minimizer support, and (v) a decreasing edit-distance profile. Genera scoring 4–5 are classified as candidate ancient taxa because they are supported by multiple independent detection and authentication criteria. Lower-scoring genera are retained in the output to allow users to inspect taxa with partial evidence.

### Simulation of ancient metagenomic data

We used gargammel (Renaud et al., 2017) to simulate ten metagenomic samples with varying human and microbial compositions, following the general simulation design of aMeta (Pochon et al., 2023). A total of 36 bacterial species were simulated, of which 18 were simulated as ancient and 18 as modern (Supplementary Table 2). Several genera (*Mycobacterium sp*., *Pseudomonas sp*., and *Streptococcus sp*.) were included in both groups to evaluate classification when ancient and modern taxa were closely related.

Species abundances in each sample were randomly assigned using a Dirichlet distribution, with a subset of taxa designated as absent. Ancient reads were simulated with damage patterns controlled by Briggs parameters (0.03, 0.4, 0.01, 0.3) (Briggs et al., 2007) and a log-normal read length distribution (loc = 3.7424, scale = 0.2795) to simulate fragmentation. Illumina sequencing errors and adapters were also incorporated, which made the final read length of 125 bp.

Each simulated sample contained approximately 500,000 ancient and 500,000 modern microbial DNA fragments, along with modern and ancient human DNA. Microbial DNA (modern + ancient) constituted 30–70% of each sample: 70%, 70%, 70%, 50%, 50%, 50%, 40%, 30%, 30%, and 30% for samples 1–10, respectively. Human DNA comprises the remaining fraction. Simulation and abundance generation scripts are available on the GitHub repository for this paper.

### Comparison of aMeta and MAT-classifier

We benchmarked MAT-classifier against aMeta using the simulated datasets. Both pipelines were run with 20 cores and 1 TB of available RAM. The Kraken2 and KrakenUniq standard bacterial databases were downloaded from https://benlangmead.github.io/aws-indexes/k2 (Langmead, 2025). Both pipelines were run using their default settings, with a 200-GB preload allocation for the KrakenUniq step in aMeta. Computational performance was evaluated from memory usage and execution time (both wall time and CPU time) using psrecord (Robitaille, 2013/2026). For MAT-classifier, trimming and Kraken2 were included when reporting the computational performance of the complete workflow.

Classification performance on simulated data was evaluated using F1 scores and receiver operating characteristic (ROC) curves. For aMeta, scores greater than 7 were used to classify ancient genera, following their recommended thresholds. For MAT-classifier, scores greater than 3 were classified as ancient, as explained above. Predicted genus classifications were compared with the ground-truth matrices to identify true positives, false positives, and false negatives, from which F1 scores were calculated:

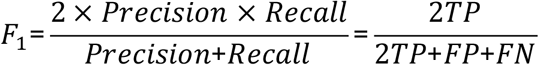

Threshold-independent performance was evaluated using ROC curves generated with the *scikit-learn* package (Pedregosa et al., 2011). Raw genus-level scores from each pipeline were compared with the ground truth across all samples, and performance was summarized using the area under the ROC curve (AUC). Unlike the F1 score, where we calculate precision and recall at predetermined thresholds, the ROC curve calculates true positive rate (TPR) and false positive rate (FPR) for all thresholds, making it threshold-independent.

### Evaluation using empirical ancient metagenomic datasets

We further compared MAT-classifier and aMeta across four types of empirical ancient datasets: (i) two deeply sequenced samples (∼60 million reads in each) from an ancient coral project (bioproject: PRJNA757238, samples: SRR16608038 and SRR16608039) (Scott et al., 2022); (ii) eight enriched 16S rRNA libraries from ancient dental calculus samples (bioproject : PRJNA1031168, samples: SRR26909287, SRR26909289, SRR26909291 to SRR26909294, SRR26909297 and SRR26909298) (Eisenhofer et al., 2024); (iii) ten shotgun libraries from ancient human dental calculus samples (bioproject: PRJNA445215, samples: SRR6877282, SRR6877284, SRR6877286, SRR6877288, SRR6877290, SRR6877310, SRR6877312, SRR6877324, SRR6877326, SRR6877393) (Mann et al., 2018); and (iv) six ancient DNA shotgun libraries from Lake Lama sediments (sedaDNA) (bioproject: PRJEB80877, samples: ERR13950600, ERR13950601, ERR13950608, ERR13950609, ERR13950613, ERR13950614) (von Hippel et al., 2025). These samples were randomly selected from big projects to limit computational requirements and were run on a cluster with 20 cores and 1 TB RAM.

## Results

### The MAT-classifier pipeline

A schematic of the MAT-classifier workflow is shown in Figure 1. The pipeline is modular, with read trimming and Kraken2 classification performed externally to provide flexibility in preprocessing and database selection. The automated workflow takes trimmed FASTQ files and Kraken2 reports as input, identifies candidate taxa based on the reports, retrieves corresponding reference genomes from NCBI, and concatenates them into genus-level reference sets. Reads are then independently aligned against each genus using either Bowtie2 or BWA-aln, with multiple alignment jobs executed in parallel.

**Figure 1.**
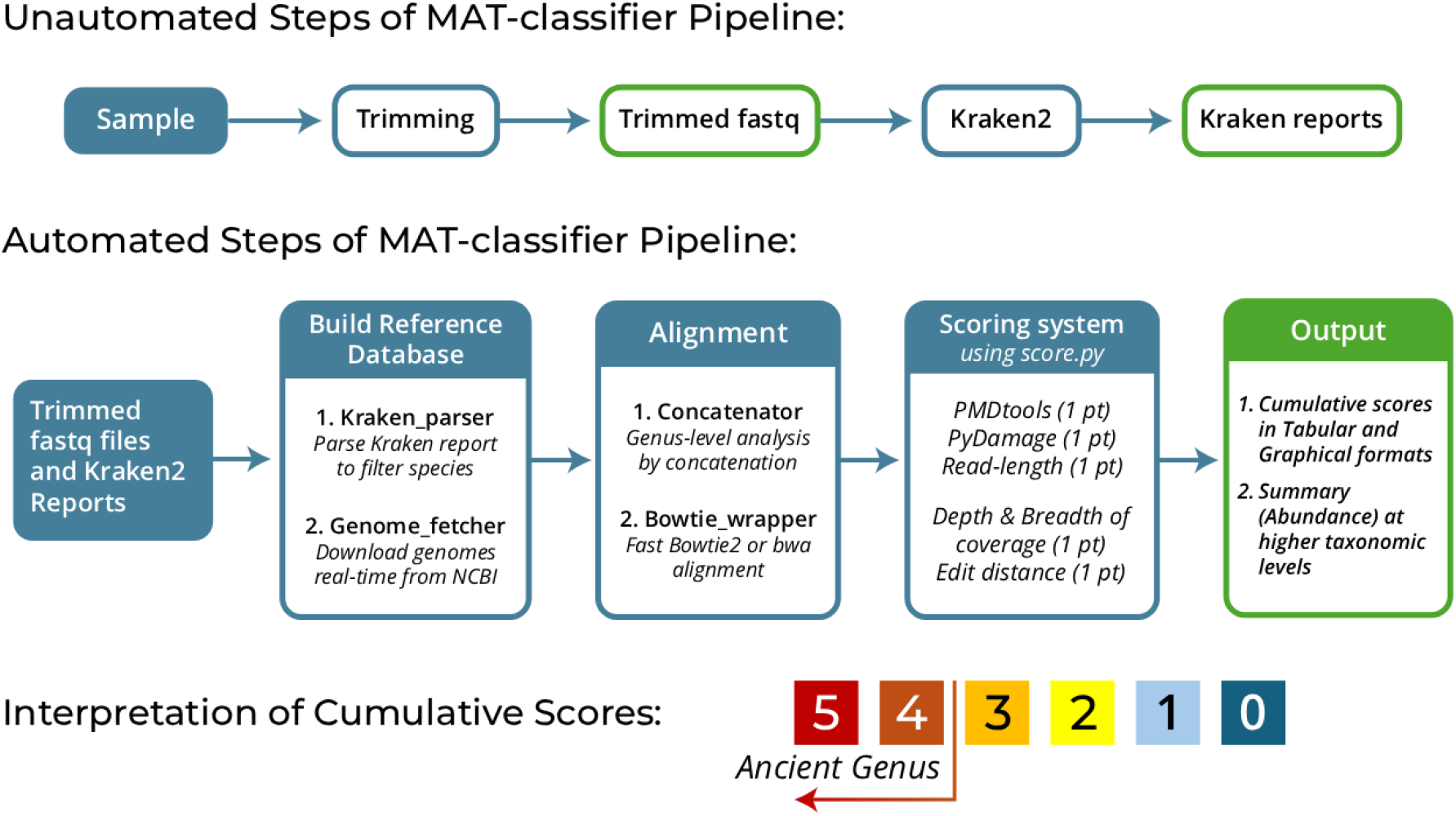
Overview of the MAT-classifier workflow. The pipeline integrates Kraken2-based candidate species selection and database creation, genus-level reference binning, independent Bowtie2 or BWA-aln alignments, and five validation and authentication scores. The output includes both tabular and graphical summaries, along with a cumulative score ranging from 0 to 5 per genus. Scores of 4–5 indicate high confidence that the genus is truly present and ancient.

The resulting alignments are evaluated using five validation and authentication criteria based on post-mortem DNA damage (pyDamage and PMDtools), fragment length, coverage breadth & depth, and edit-distance profiles. These criteria are integrated into a cumulative score ranging from 0 to 5 for each genus, with scores of 4–5 indicating candidate ancient taxa.

### Computational Performance of MAT-classifier against simulated data

We benchmarked MAT-classifier against aMeta, a recently developed ancient metagenomic workflow with a similar hybrid classification strategy for efficiency. Both pipelines were evaluated on 10 simulated ancient metagenomic datasets using 20 CPU cores and up to 1 TB of RAM. MAT-classifier was tested with both Bowtie2 and BWA-aln to assess whether aligner choice affected computational performance.

MAT-classifier showed substantially lower memory usage and shorter runtimes than aMeta with either aligner (Figure 2, Supplementary Figure 2). During the k-mer-based classification step, Kraken2 used approximately 97 GB of peak memory, whereas the KrakenUniq stage of aMeta approached the available 1-TB memory limit despite a preload size of 200 GB (Figure 2.ii). This difference may be reduced on systems with limited memory. However, the difference was more pronounced during the downstream stage, which includes alignment: aMeta repeatedly used approximately 140 GB of memory during MALT alignment, whereas MAT-classifier required approximately 7 GB at peak (Figure 2.iii). Unlike the k-mer-based classification step, memory usage during the alignment stage cannot be reduced. Moreover, MAT-classifier’s memory usage can be further reduced by using a size-capped Kraken2 database while still retaining an accuracy advantage over aMeta (Supplementary Figures 3 and 4).

**Figure 2.**
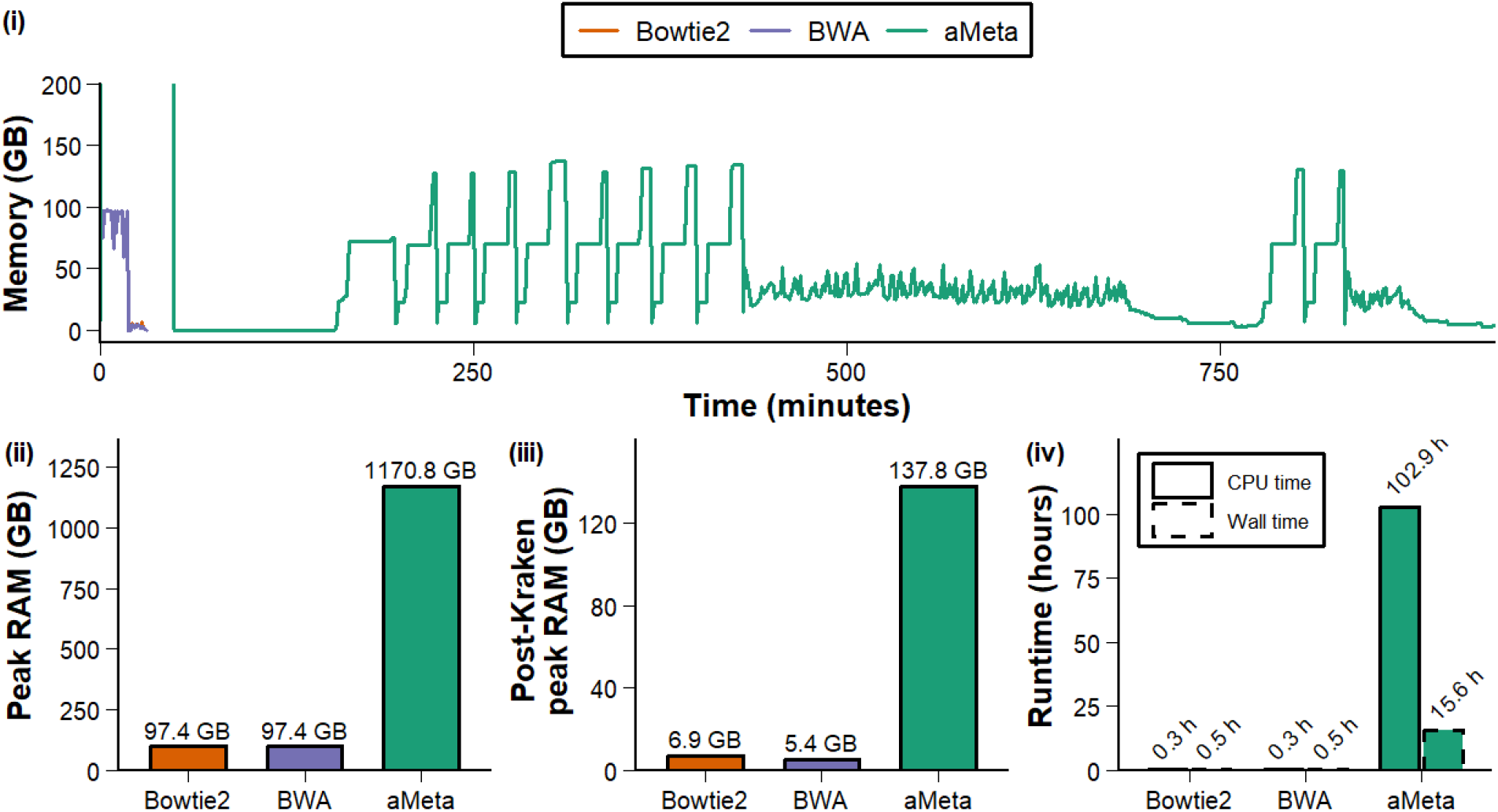
Computational performance of MAT-classifier and aMeta on simulated ancient samples. (i) Memory usage over time for MAT-classifier using Bowtie2 (orange), MAT-classifier using BWA (violet), and aMeta (green). The y-axis is restricted to 0–200 GB to improve visualization, as aMeta exceeds this range during the initial hour. (Full figure in Supplementary Figure 2) (ii) Peak memory usage of the entire pipeline and (iii) peak memory usage excluding k-mer-based classification. The peak memory after Kraken essentially indicates memory consumption during alignment. (iii) Total CPU time and wall-clock time for MAT-classifier and aMeta. MAT-classifier estimates include the external preprocessing steps of read trimming and Kraken2 classification.

Runtime differences were similarly substantial. MAT-classifier completed the full workflow in approximately 30 minutes, whereas aMeta required approximately 15 hours under the same computational allocation (Figure 2.iv). Aligner choice had little effect on MAT-classifier resource usage. Bowtie2 and BWA-aln showed similar memory profiles and runtimes. Hence, the subsequent benchmarking on empirical datasets was performed using Bowtie2 only.

### Classification accuracy on simulated data

We next evaluated the ability of each pipeline to distinguish ancient taxa from modern or absent taxa in the simulated datasets. Because aMeta and MAT-classifier use different scoring scales (0–10 and 0–5, respectively), we compared their raw scores using receiver operating characteristic (ROC) curves, which evaluate classification performance across all possible score thresholds.

MAT-classifier produced a higher area under the ROC curve (AUC) than aMeta, with an AUC of 0.87 compared with 0.79 for aMeta (Figure 3). This indicates that MAT-classifier consistently assigned higher scores to true ancient genera than to modern or absent genera across the simulated dataset. MAT-classifier produced similar ROC performance with Bowtie2 and BWA-aln, suggesting that aligner choice had little effect on classification accuracy under the simulated conditions.

**Figure 3:**
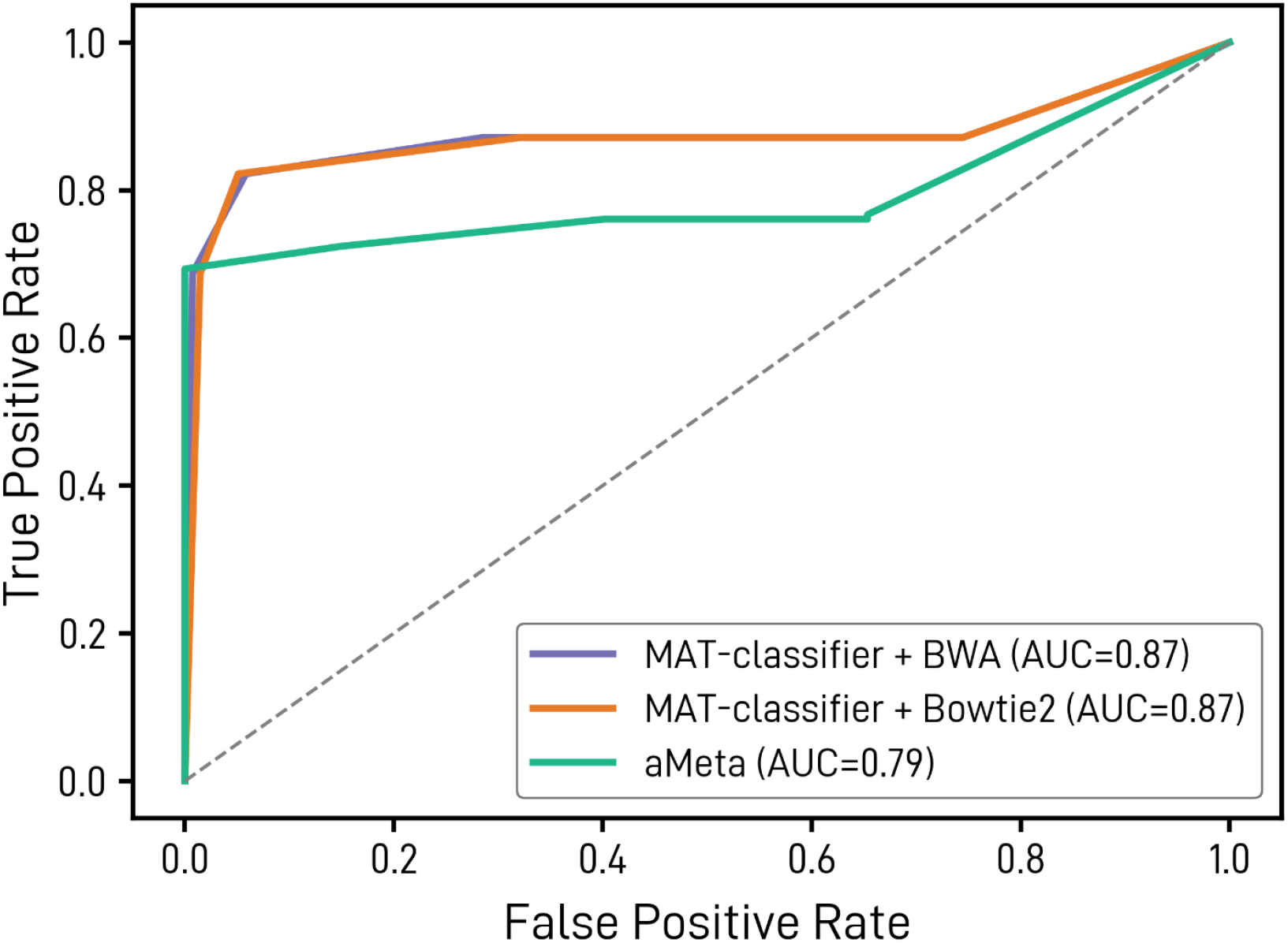
Receiver operating characteristic (ROC) curves comparing MAT-classifier and aMeta on simulated datasets. ROC curves were generated from raw genus-level scores across all simulated samples for MAT-classifier using Bowtie2 (orange), MAT-classifier using BWA-aln (violet), and aMeta (green). AUC values are shown in the legend.

### Accuracy at recommended cutoffs

Because users ultimately classify taxa using discrete score thresholds, we compared pipeline performance at the recommended cutoffs: >7 for aMeta and >3 for MAT-classifier. The MAT-classifier threshold was selected because genera scoring >3 must be supported by multiple validation criteria and at least one authentication criterion.

Accuracy was evaluated using F1 scores, which balance precision and recall. For each classifier, taxa exceeding the respective thresholds were classified as ancient. True positives were ancient genera present in the simulated ground truth, whereas false positives included absent or modern genera incorrectly classified as ancient. MAT-classifier achieved significantly higher F1 scores than aMeta across samples (one-sided Wilcoxon signed-rank test, *p* = 0.0010; Figure 4), demonstrating improved accuracy at the recommended cutoff.

**Figure 4:**
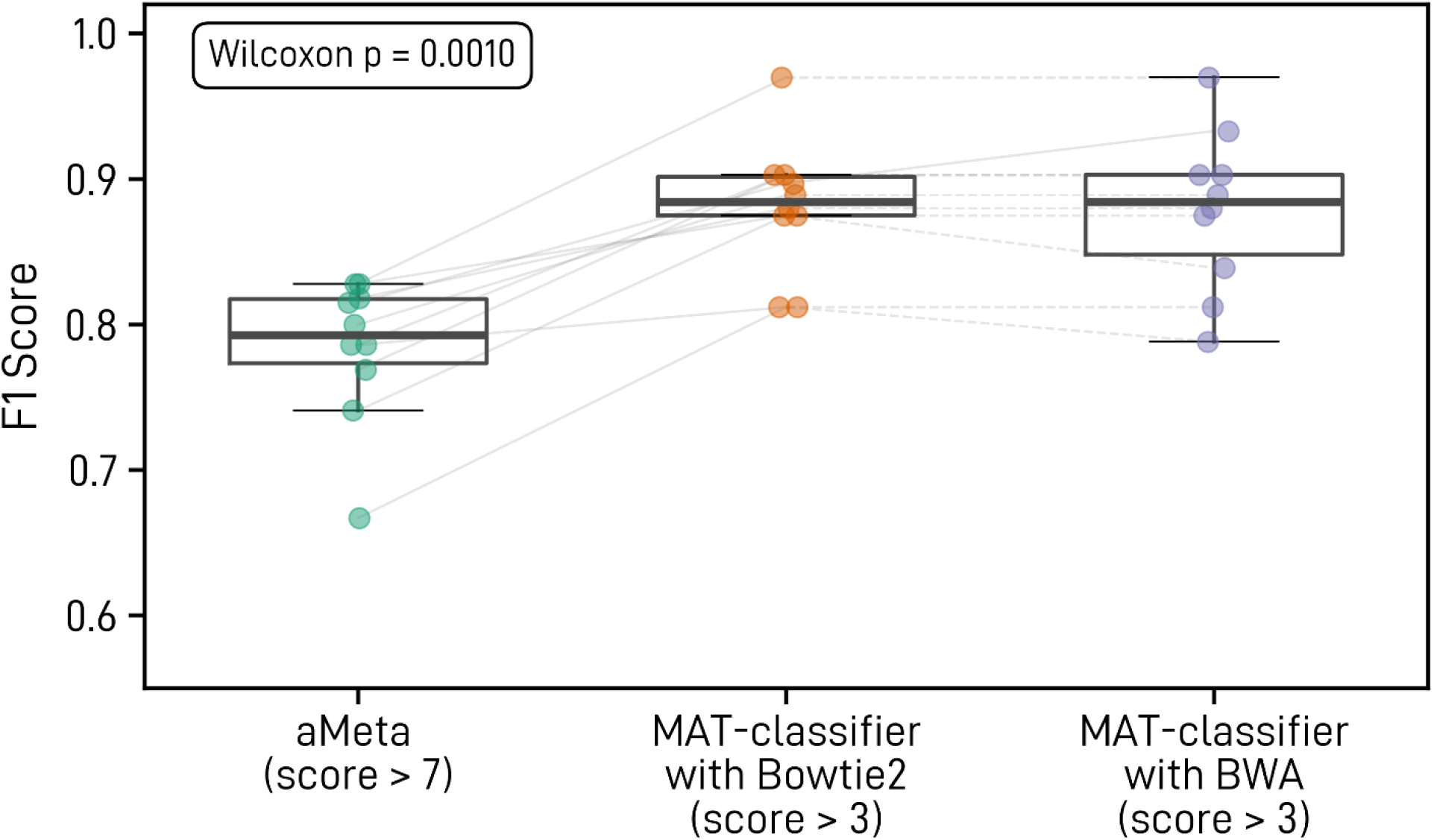
Ancient-taxon classification accuracy at recommended score cutoffs. F1 scores are shown for aMeta (green, score > 7), MAT-classifier with Bowtie2 (orange, score > 3), and MAT-classifier with BWA (violet, score > 3). Each point represents one simulated sample, with paired results connected by grey lines. MAT-classifier achieved higher F1 scores than aMeta across all samples (one-sided Wilcoxon signed-rank test, *p* = 0.0010).

Notably, Bowtie2 and BWA-aln produced similar F1 scores for most samples. Bowtie2 performed better for samples 1 and 9, whereas BWA-aln performed better for sample 6.

These differences resulted primarily from one aligner failing to detect low-abundance ancient genera, suggesting that aligner choice had only a minor effect on classification performance.

We also evaluated performance when ancient and modern species from the same genus co-occurred. The simulations included *Mycobacterium* (one ancient and one modern species), *Pseudomonas* (one ancient and three modern species), and *Streptococcus* (one ancient and two modern species). MAT-classifier detected the ancient genus in 6 of 7 *Mycobacterium*, 7 of 9 *Pseudomonas*, and 5 of 8 *Streptococcus* occurrences. In comparison, aMeta failed to identify ancient *Pseudomonas* in all of these cases but detected two additional *Streptococcus* occurrences relative to MAT-classifier. Overall, these results indicate that Mat-classifier can recover ancient signals reliably even when modern species from the same genus are present, although sensitivity decreases for low-abundance ancient taxa.

### Computational performance on real-world datasets

We further benchmarked the MAT-classifier against aMeta using four empirical ancient metagenomic datasets: (i) shotgun libraries from ancient corals (Scott et al., 2022), (ii) enriched 16S rRNA libraries from ancient dental calculus samples (Eisenhofer et al., 2024); shotgun libraries from ancient human dental calculus samples (Mann et al., 2018); and shotgun libraries from Lake Lama sediments (sedaDNA) (von Hippel et al., 2025). All analyses were done using 40 cores and 1 TB RAM.

Consistent with the simulation results, MAT-classifier showed lower wall-clock time and memory usage than aMeta across the empirical datasets (Figure 5). However, MAT-classifier took similar or higher CPU time (Figure 5.i). This indicates that MAT-classifiers make more effective use of parallel execution, reducing elapsed runtime by prioritizing computational power.

**Figure 5:**
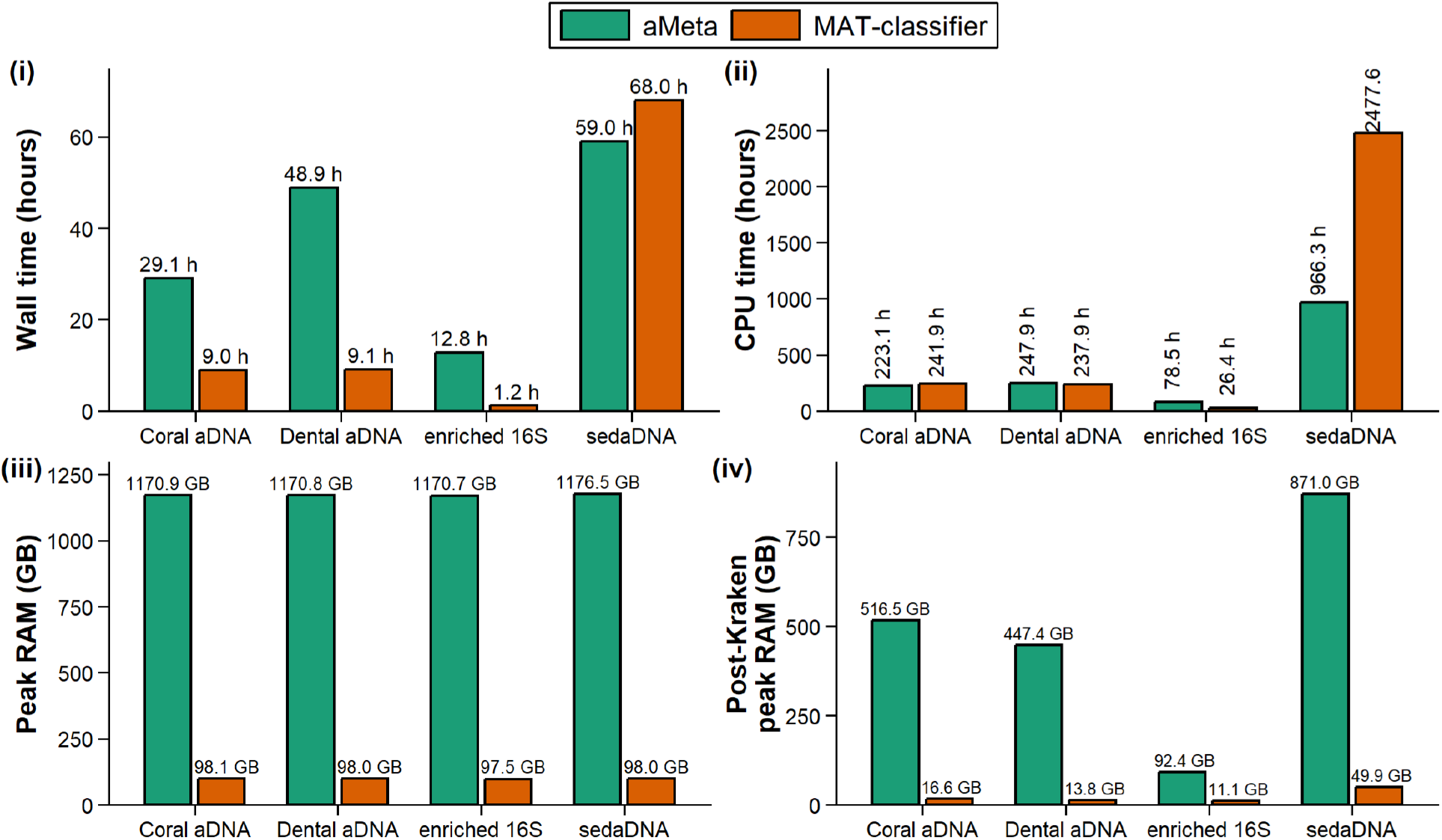
Computational performance of MAT-classifier and aMeta across empirical ancient metagenomic datasets. MAT-classifier using Bowtie2 (orange) and aMeta (green) were compared across four datasets using four metrics: (i) Wall-clock time, (ii) total CPU time, (iii) Peak RAM usage for the complete workflows, and (iv) Peak RAM usage excluding the Kraken2 and KrakenUniq classification steps, highlighting differences in memory requirements during downstream processing.

MAT-classifier also required substantially less memory than aMeta across datasets (Figure 5.iii and 5.iv). This difference remained even when the Kraken2 and KrakenUniq classification steps were excluded from the comparison (Figure 5.iv), indicating a massive reduction in memory usage during the alignment process. This result highlights the effectiveness of MAT-classifier’s unique design, which helps reduce required memory during the alignment stage.

Across the empirical datasets, MAT-classifier identified substantially more candidate ancient genera than aMeta (Supplementary Table 3). MAT-classifier recovered 652, 108, 173, and 97 genera not detected by aMeta in the sedaDNA, enriched 16S, dental aDNA, and coral aDNA datasets, respectively. Importantly, MAT-classifier was able to recover most genera reported by those studies (see Supplementary Table 3). For example, it detected *Ruegeria*, the principal genus reported from the ancient coral dataset, whereas aMeta didn’t. Agreement was lowest for the enriched 16S dataset, probably because MAT-classifier is more suitable for metagenomic data, although the original study didn’t authenticate reported taxa using damage patterns, limiting direct comparison.

## Discussion

Ancient microbiome research is still a comparatively young branch of palaeogenomics, and computational approaches for taxonomic detection and authentication remain under active development. Existing alignment-based workflows can require substantial computational resources. To address this limitation, we developed MAT-classifier with an emphasis on reducing memory requirements. Rather than performing large-scale competitive metagenomic alignment, MAT-classifier first restricts the candidate taxonomic space using Kraken2 and subsequently performs independent alignments against smaller genus-level reference sets using traditional aligners like Bowtie2 or BWA-aln.

Benchmarking with both simulated and empirical datasets highlights MAT-classifier’s lower memory consumption. Moreover, in simulated datasets, MAT-classifier achieved higher accuracy, indicating improved detection of ancient taxa under these conditions. Bowtie2 and BWA-aln produced broadly similar computational and classification performance, suggesting that users can select an aligner according to their needs without substantially altering overall pipeline performance. Together, these results support MAT-classifier as a computationally accessible approach for genus-level detection and authentication of ancient microbial DNA.

### Memory efficiency through genus-level alignment and Kraken2 screening

The lower memory requirement of MAT-classifier primarily results from dividing the alignment problem into smaller genus-level reference sets rather than loading and searching a large metagenomic reference database. Independent alignment jobs are distributed across available CPU cores, taking advantage of parallel computation while keeping the memory requirement of each individual alignment relatively small.

Kraken2 screening further reduces the downstream alignment burden by restricting reference construction to plausible candidate taxa. Although Kraken2 consumes the most memory in the whole pipeline, its resource requirements can be adjusted according to the user’s needs. For example, smaller Kraken2 databases can be generated using database-size restrictions. Alternatively, KrakenUniq with preload size restriction can be used to reduce RAM requirements at the expense of increased runtime. We therefore made the Kraken2 classification step external for flexibility.

### Trade-offs of independent mapping

Implementing traditional aligners over competitive metagenomic aligners means MAT-classifier has to trade large memory requirements for greater dependence on computational power. Metagenomic aligners such as MALT search many reference genomes simultaneously but require substantial memory, whereas MAT-classifier performs smaller independent alignments requiring more computational power but less memory. To achieve efficiency, our pipeline distributes the alignment jobs across available CPU cores. Consistent with this design, MAT-classifier showed similar or higher CPU usage to aMeta while requiring substantially less wall-clock time and memory. On systems with fewer CPU cores, runtimes may increase, but the lower memory requirement should make the workflow more practical on modest workstations and computing nodes.

Independent mapping also increases the risk of cross-mapping because the same read may align separately to multiple related reference genomes. MAT-classifier limits this problem by constructing genus-level reference sets, thereby reducing redundant species-level assignments. Also, spurious alignments are penalized with scoring criteria that check for sufficient taxonomic depth and breadth. Nevertheless, genus-level binning cannot completely eliminate cross-mapping between closely related genera. Hence, we recommend users to proceed with caution when interpreting closely related taxa.

### Accuracy and limitations of MAT-classifier

MAT-classifier showed higher classification accuracy than aMeta in our simulated benchmarks. Several features may contribute to this performance. Our design of Genus-level consolidation may increase the statistical power by pooling evidence. In addition, the permissive Kraken2 screening thresholds (≥100 reads and ≥500 distinct minimizers) retain a broad set of candidate taxa, whereas a more stringent validation threshold (≥200 reads and ≥1,000 distinct minimizers) and multiple authentication criteria during the scoring stage reduce false-positive classifications.

MAT-classifier nevertheless has important limitations. Firstly, the workflow restricts classifications at the genus level rather than attempting species-level identification. This design is deliberate, as short ancient reads and high sequence similarity among closely related microbial species can make species-level assignments unreliable, especially due to microbial taxonomy’s continuing revision (Hleap et al., 2021) and differences in taxonomic databases (Duvallet et al., 2017).

In addition, MAT-classifier identified substantially more candidate taxa than aMeta in empirical datasets. Although this may reflect increased sensitivity, some of these additional calls may be false positives. We therefore recommend using MAT-classifier scores as a screening framework rather than as sole evidence of ancient origin, with important or unexpected taxa further evaluated using their underlying authentication evidence, particularly terminal post-mortem damage patterns plotted by PyDamage. MAT-classifier also lacks a dedicated pathogen-screening module, although its modular design allows flexible modifications which can be implemented using the Kraken2 classifications.

## Conclusions

MAT-classifier provides an accurate and computationally efficient framework for genus-level classification and authentication of ancient microbial DNA. As demonstrated using simulated and empirical datasets, its low memory requirements and multi-metric validation approach make ancient microbiome analysis more accessible across a wider range of computational environments.

## Declarations

### Availability of data and materials

The pipeline is publicly available at https://github.com/AranyaDhibar/MAT-classifier. The scripts for simulated metagenomic datasets and other scripts used for this article can be accessed at https://github.com/AranyaDhibar/scripts_matclassifier_paper. The simulated datasets, runtime datasets, as well as the snapshots of the above-mentioned GitHub repository, are archived in the Zenodo digital repository (https://doi.org/10.5281/zenodo.22822502).

### Author Contributions

A.D. and M.V.M. conceptualized the study. A.D. wrote the scripts, performed the analysis, and wrote the draft manuscript under the supervision of M.V.M. All authors edited the manuscript and approved the final version.

## Acknowledgements

This work was supported by the National Science Foundation grants OCE-2318775 to M. V. M. All required computational analyses were carried out using the computational resources of the Texas Advanced Computing Center (TACC) at The University of Texas at Austin. URL: http://www.tacc.utexas.edu.

