## Supplemental files for "MAT-classifier: A memory-efficient pipeline for accurate genus-level profiling from ancient metagenomic data"

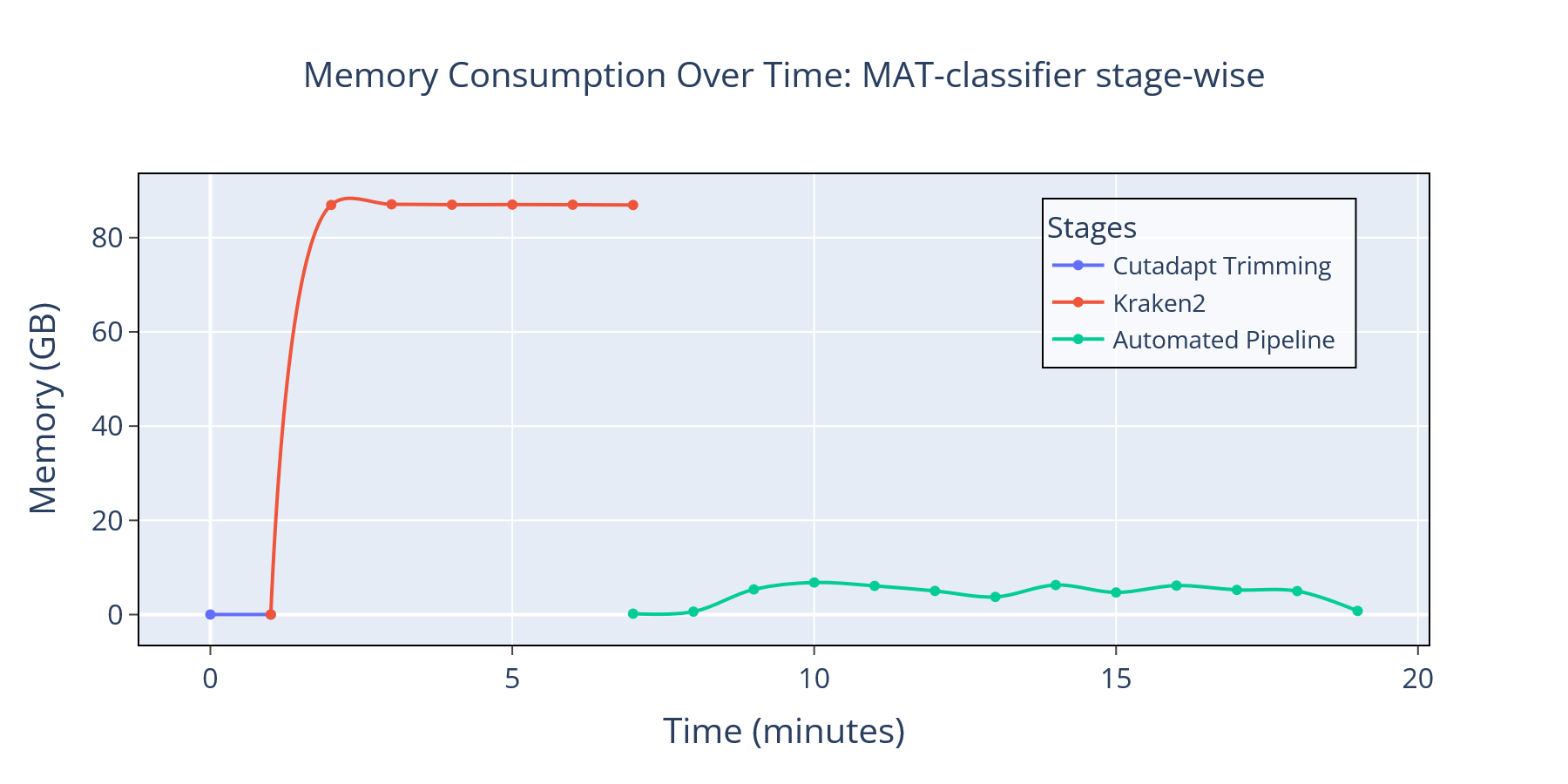

**Supplementary figure 1:** Memory consumption over time plot for the MAT-classifier pipeline color coded according to different steps – external trimming step (blue), external Kraken2 step (orange) and the automated pipeline (green). This plot highlights Kraken2 is the most memory intensive step for the whole pipeline with peak usage of almost 90 GB memory. Whereas the automated part which performs Bowtie2 alignment and other operations consumes very little memory with peak usage of 7 GB.

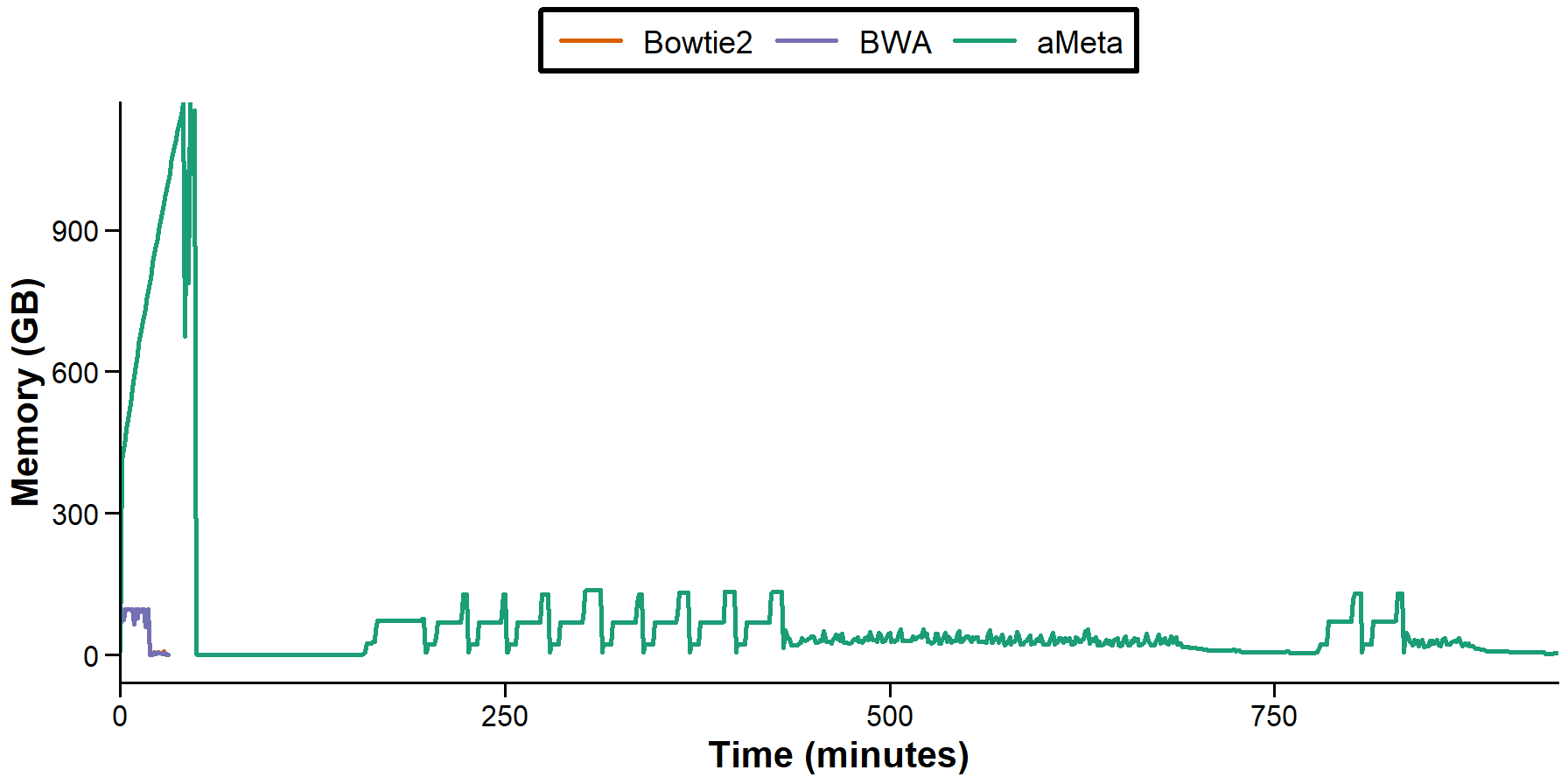

**Supplementary figure 2:** The full version of memory usage over time plot for MAT-classifier using Bowtie2 (orange), MAT-classifier using BWA (violet), and aMeta (green).

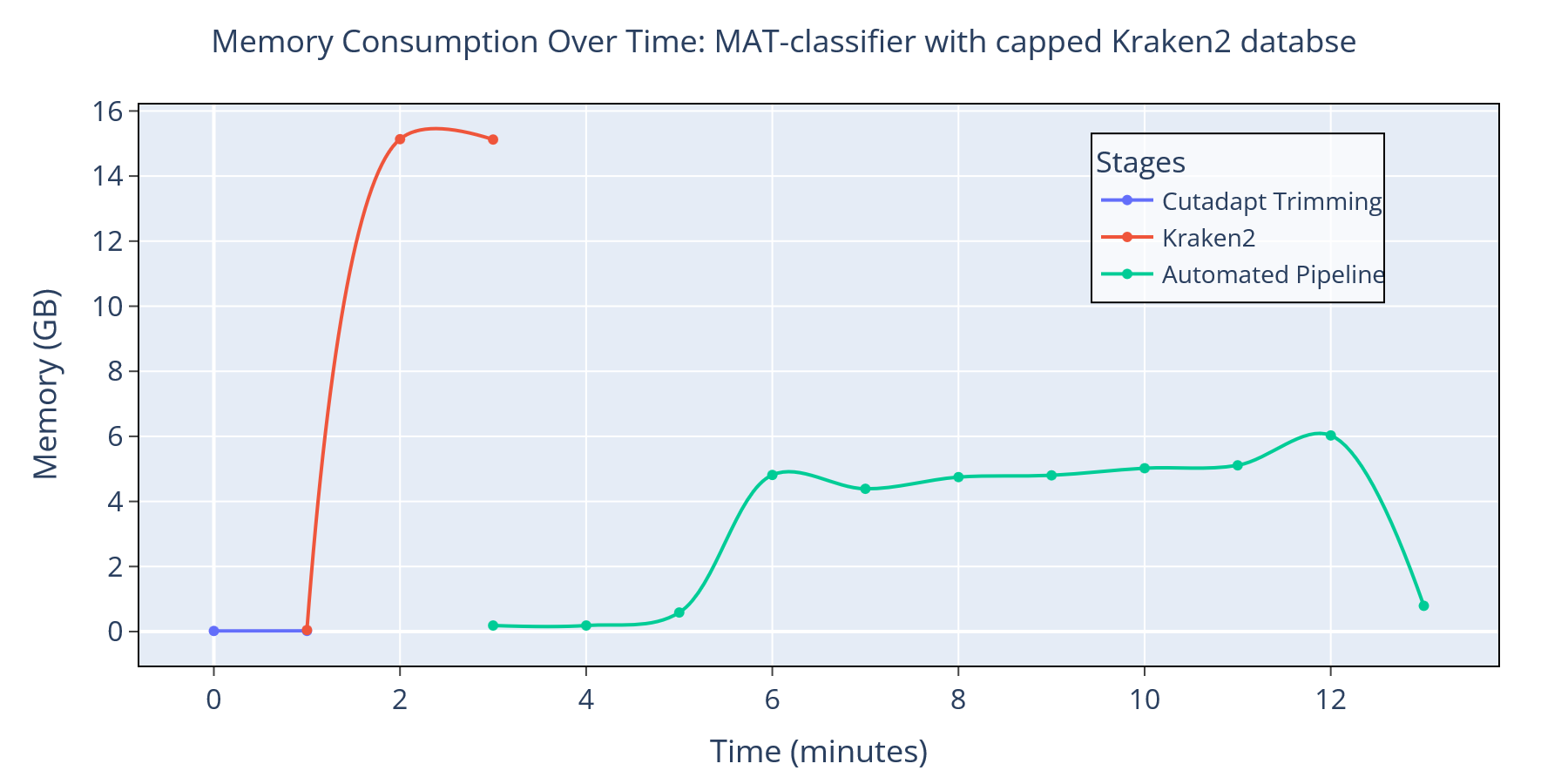

**Supplementary figure 3:** Memory consumption over time plot for the MAT-classifier pipeline with 16GB capped Kraken2 database color coded according to different steps – external trimming step (blue), external Kraken2 step (orange) and the automated pipeline (green). Peak memory usage was reduced to ~16 GB.

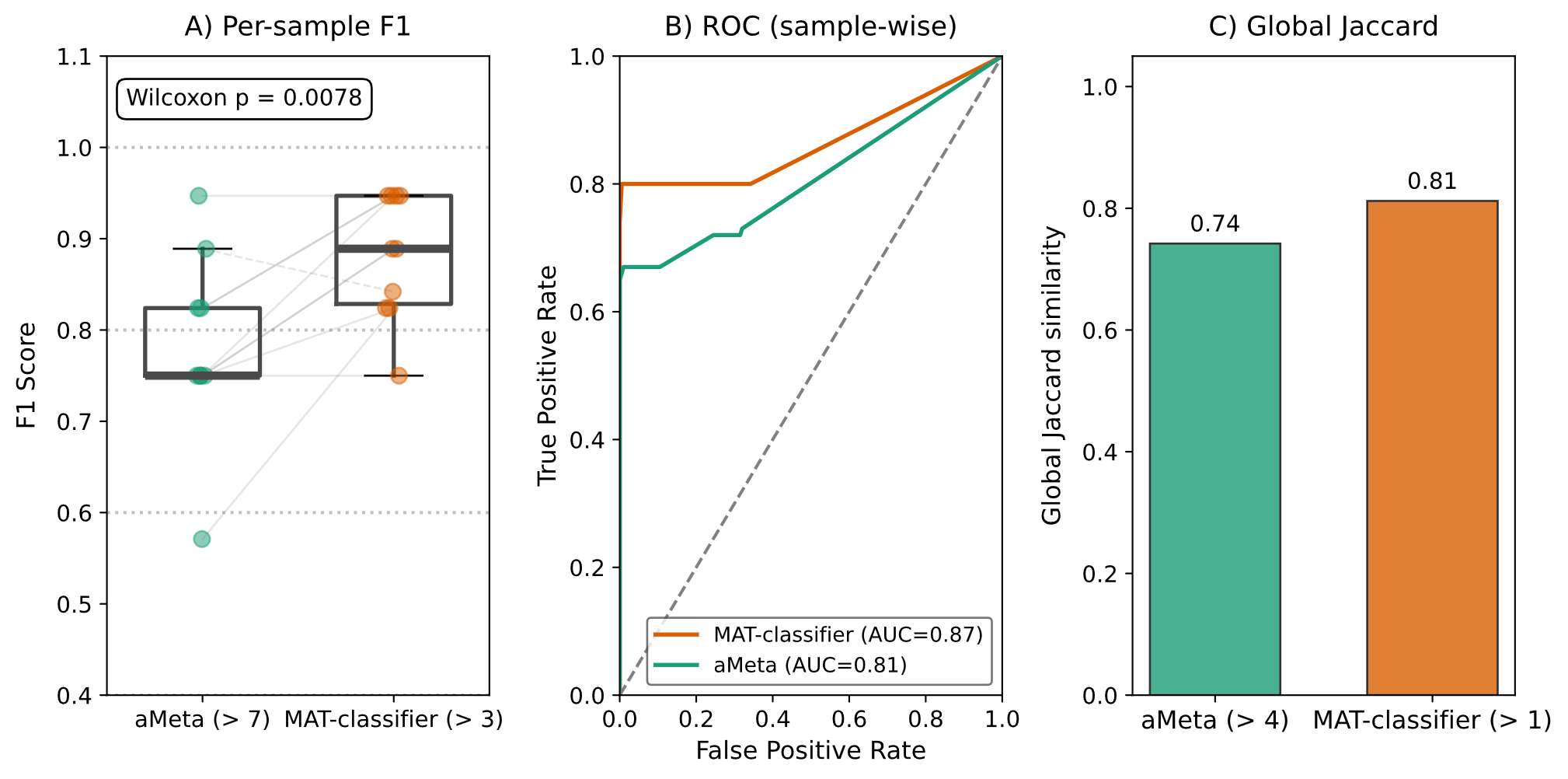

**Supplementary figure 4:** Performance of MAT-classifier with a 16 GB capped Kraken2 database compared with aMeta. The simulated dataset version 1 was used here. (A) Sample-wise F1 scores (aMeta >7, green; MAT-classifier >3, orange). Paired samples are connected by grey lines (solid lines = increase, dotted lines = decrease). In 9 of 10 samples, MAT-classifier achieved higher F1 scores, with significant improvement over aMeta (Wilcoxon signed-rank test, *p* = 0.0078). (B) ROC curves of sample-wise scores showing higher AUC for MAT-classifier, indicating stronger discriminative ability even with a capped database. (C) Global Jaccard similarity for taxa detection (ancient + modern taxa), showing retained improvements of MAT-classifier relative to aMeta.

| Reads cutoff | Unique k-mers cutoff | AUC (sample-wise) |
| --- | --- | --- |
| 50 | 250 | 0.91320 |
| 75 | 375 | 0.91423 |
| 100 | 500 | 0.91723 |
| 125 | 625 | 0.91720 |
| 150 | 750 | 0.91720 |

**Supplementary table 1:** AUC values calculated for different filter thresholds during database construction based on Kraken reports. The AUC was calculated using the final scores and ground truth. Higher AUC means higher accuracy. Smaller cutoffs increase the database size but reduce sensitivity–specificity balance. Cutoffs of ≥100 reads and ≥500 distinct k-mers yielded the highest discriminatory performance. We therefore selected the 100/500 cutoff to maximize the number of candidate species for database construction, while maintaining high discriminatory performance.

| **Ancient species** | **Modern species** |
| --- | --- |
| Bradyrhizobium erythrophlei | Acinetobacter baumannii |
| Campylobacter rectus | Bacteroides fragilis |
| Clostridium botulinum | Brevibacterium aurantiacum |
| Enterococcus faecalis | Burkholderia mallei |
| Mycobacterium avium | Cutibacterium acnes |
| Mycolicibacterium aurum | Lactococcus lactis |
| Neisseria meningitidis | Methylobacterium bullatum |
| Nocardia brasiliensis | Micromonospora echinospora |
| Parvimonas micra | Mycobacterium riyadhense |
| Prosthecobacter vanneervenii | Nonomuraea gerenzanensis |
| Pseudomonas thivervalensis | Planobispora rosea |
| Ralstonia solanacearum | Pseudomonas aeruginosa |
| Rothia dentocariosa | Pseudomonas caeni |
| Salmonella enterica | Pseudomonas psychrophila |
| Sorangium cellulosum | Rhodococcus chondri |
| Streptococcus pyogenes | Rhodococcus hoagii |
| Streptosporangium roseum | Streptococcus pneumoniae |
| Yersinia pestis | Streptococcus pyogenes |

**Supplementary table 2:** List of species simulated as Ancient or modern species in the simulated samples. Blue colored cells highlight taxa present in both modern and ancient datasets

| **Dataset** | **Taxa predicted by MAT-classifier** | **Taxa predicted by aMeta** | **Common to MAT-classifier and aMeta** | **Comparison with original study** |
| --- | --- | --- | --- | --- |
| SedaDNA | 652 | 0 | 0 | MAT-classifier recovered 4 genera out of 6 ancient taxa reported in the original study from the selected samples, including low-abundance taxa such as *Delftia.* However, aMeta didn’t recover any ancient taxa. |
| Enriched 16S | 114 | 10 | 6 | The original study reported taxa without ancient-DNA damage authentication, limiting direct comparison. MAT-classifier only recovered 24 taxa out of 57 reported in the study, whereas aMeta recovered none of them. This dataset may also be less suitable for MAT-classifier and aMeta because the workflows were designed primarily for shotgun metagenomic data rather than 16S libraries. |
| Dental aDNA | 187 | 14 | 14 | MAT-classifier recovered 14 out of the top 16 genera reported in the original study. On the other hand, aMeta recovered 6 out of the 16 genera. |
| Coral aDNA | 103 | 6 | 6 | MAT-classifier recovered *Ruegeria*, the genus confidently reported in the original study, whereas aMeta did not. Other microbial classifications in the original study were primarily reported at the phylum level and were therefore not compared. |

**Supplementary table 3:** Comparison of ancient genera identified by MAT-classifier and aMeta across empirical datasets. Counts indicate number of genera classified as ancient by either pipelines and the common set between them. Recovery of taxa reported in the original studies is summarized for additional context; these comparisons should not be interpreted as ground truth because authentication criteria differed among studies.
